# *Nissolia brasiliensis* as a non-nodulating model legume

**DOI:** 10.1101/2025.07.08.663655

**Authors:** Camille Girou, Jean Keller, Cyril Libourel, Fabian van Beveren, Matheus Bianconi, Isabelle Dufau, Charlotte Cravero, Caroline Callot, Nathalie Rodde, Olivier Coriton, Virginie Huteau, Pierre-Marc Delaux, Tatiana Vernié

## Abstract

The nitrogen-fixing root nodule symbiosis (RNS) is specifically formed by four orders of angiosperms. The largest of these four orders include the legume family, the Fabaceae. Among legumes, historical model species have emerged, such as the RNS-forming *Medicago truncatula* and *Lotus japonicus*, or, more recently, *Aeschynomene aevenia*. By contrast, legume species that have lost RNS have been largely ignored. Here, we describe the first chromosome-level assembly for a non-RNS-forming legume, the tropical papilionoid *Nissolia brasiliensis*. We compared its genome to closely related legumes and identified genes associated with RNS. Finally, we developed a stable transformation protocol that can be deployed in the future to re-evolve RNS in legumes, a first step toward the goal of engineering RNS in non-legume crops.

## Introduction

Nitrogen-fixing mutualistic symbioses where a diazotrophic bacteria provides fixed-nitrogen to a eukaryotic host have evolved multiple times across the tree of life (Xu and Wang, 2023). Such symbiotic functionality has repeatedly emergedin the plant kingdom, with the level of interaction ranging from loose association on the plant surface (Van Deynze et al., 2018), to endophytism in cavities (Li et al., 2018) canals (Li et al., 2020) or roots (Liu et al., 2022). The closest level of association has been achieved in two clades of flowering plants: the genus *Gunnera,* where the nitrogen-fixing cyanobacteria are hosted inside plant cells from the internal layer of stem glands (Bergman et al., 1992; Parniske, 2018) and four monophyletic orders (Fagales, Fabales, Cucurbitales and Rosales) collectively referred as the Nitrogen-Fixing root Nodule (NFN) clade (Kistner and Parniske, 2002). In the nitrogen-fixing root nodule symbiosis (RNS), bacteria from either the rhizobia assemblage or the *Frankia* genus colonize dedicated plant organs, the nodule, whose formation is triggered by the host plants following the perception of the symbiont (Kistner and Parniske, 2002). The colonisation results in most cases in the trans- or intra-cellular accommodation of the symbiont (Kundu et al., 2025).

RNS has been particularly studied as the trait is found in crop legumes, such as soybean and beans. A singularity of the RNS is its high loss rate along the evolution of the NFN clade, with multiple species not being able to engage in this symbiosis (Griesmann et al., 2018; Van Velzen et al., 2018). In contrast with other clades, this loss rate is much lower in legumes from the Papilionoids subclade, presumably because of the evolution of an additional symbiotic stage, the symbiosome, that provides higher control of the host upon the symbionts and thus possibly limits the emergence of cheaters (De Faria et al., 2022). Several model species have been developed by the scientific community in papilionoid legumes. Besides the historical models *Medicago truncatula* and *Lotus japonicus*, other species have recently emerged to study specific symbiotic features. This is the case of *Aeschynomene aevenia* (Quilbé et al., 2021), which is able to form RNS independently of the canonical symbiotic Nod-factor signal; *Arachis hypogeia,* which is infected by the rhizobial symbiont via a non-conventional crack-entry mode (Karmakar et al. 2019); or *Lupinus albus,* which forms atypical nodules, and has lost the ancestral arbuscular mycorrhizal symbiosis (Hufnagel et al., 2020).

Among the papilionoid legume, the *Nissolia* genus was proposed as non-nodulating based on multiple root sampling in the wild (Sprent, 2007). Furthermore, the draft sequence of *Nissolia schottii*, a species native to the USA and Mexico, revealed the absence of *NIN* and *RPG,* two genes essential for RNS, a feature shared with most other species from the Fabales, Fagales, Cucurbitales and Rosales which have lost RNS (Griesmann et al., 2018; Van Velzen et al., 2018). A PCR-based approach suggested the absence of *NIN* in two other *Nissolia* species, *N. scandens* and *N. brasiliensis* (formerly *Chaetocalyx scandens* and *C. brasiliensis*, (Moura et al., 2018) and thus the likely loss of RNS before the diversification of the *Nissolia* genus (Griesmann et al., 2018). The genus *Nissolia* belongs to the Dalbergioid clade and is composed of 29 species, all endemic to the Americas, with the distribution ranging from Argentina to Mexico and the South of the USA (Moura et al., 2018). Because of its clear non-RNS-host status the *Nissolia* genus represents the ideal context to develop a non-RNS-forming model legume species. Such model would facilitate the study of RNS using comparative biology approaches, but also provide a background for re-evolving RNS using synthetic biology, hence bringing back a trait once existing in the lineage. However, only a fragmentary genome assembly of *N. schottii* sequenced using short-read sequencing technology (Griesmann et al., 2018) is currently available for the *Nissolia* genus. In addition, stable genetic transformation has never been achieved, strongly limiting its deployment as a model system.

Here, we describe a long read-based genome assembly of *Nissolia brasiliensis*, and use a comparative phylogenomic approach to identify previously overlooked RNS-associated genes. To promote the experimental manipulation of the species, we develop a stable genetic transformation protocol, providing a useful resource that makes *N. brasiliensis* a suitable non-RNS forming model legume.

## Results

### The Nissolia brasiliensis genome

The genus *Nissolia* as a whole has been reported as non-RNS forming, but it is supposed to have maintained the more ancient arbuscular mycorrhizal symbiosis (AMS). To test these hypotheses, we grew the species *N. brasiliensis* in the presence of the AM fungus *Rhizophagus irregularis,* and of five different rhizobial strains in parallel. While nodules were not observed in any of the tested conditions (Supplementary Table S1), consistent mycorrhizal colonisation was scored in 14/15 plants inoculated with *R. irregularis* (Supplementary Figure S1). This initial test further supported the selection of *N. brasiliensis* as a potential model for non-RNS forming papilionoids.

DNA was extracted from young leaves, and sequenced on a PacBio HiFi platform, yielding a total of 1,647,178 corrected reads with a N50 of 18.1Mbp. Genome assembly resulted in a total of 647 scaffolds with a N50 of 35.7Mb and a BUSCO completion score of 93.1% against the Fabales dataset (n=7843) (Table 1&2). The assembly total size was ∼ 790Mb, close to the average 863Mb estimated by flux cytometry, which also identified 20 chromosomes (10 pairs; Supp. Table S2 and Supp. Figure S2). The annotation of *N. brasiliensis* genome has been performed based on RNA-seq and protein hints, and predicted a total of 26,850 protein-coding genes, resulting in a BUSCO completion of 92.7% (Table 2). Comparison with the model legume *M. truncatula* revealed important synteny between the two species based on predicted gene models (Supp. Figure S3).

**Table 1.** Assembly statistics of the Nissolia brasiliensis genome.

|  |  | NGS | Optical Map | Hybrid Scaffold | Hybrid Scaffold and NGS sequences not scaffolded assembly |
| --- | --- | --- | --- | --- | --- |
| Raw data (Gb) |  | 433 | 437 |  |  |
| Assembly | Count | 654 | 137 | 37 | 647 |
|  | N50 length (Mb) | 37,492 | 28,27 | 35,705 | 35,705 |
|  | Max length (Mb) | 91,028 | 50,628 | 90,644 | 90,644 |
|  | Total length (Mb) | 776,339 | 915,542 | 752,456 | 790,641 |

**Table 2.** Assembly completeness of the Nissolia brasiliensis genome evaluated by BUSCO score on Fabales odb10 database (n=7843)

| Categories | Hybrid Scaffold and NGS sequences not scaffolded assembly |  | Annotated genome |  |
| --- | --- | --- | --- | --- |
|  | BUSCOs | % | BUSCOs | % |
| Complete BUSCOs | 7304 | 93.1 | 7268 | 92.7 |
| Complete and single-copy BUSCOs | 6602 | 84.2 | 6492 | 82.8 |
| Complete and duplicated BUSCOs | 702 | 9 | 776 | 9.9 |
| Fragmented BUSCOs | 95 | 1.2 | 63 | 0.8 |
| Missing BUSCOs | 444 | 5.7 | 512 | 6.5 |

To place the *N. brasiliensis* genome in an evolutionary context within the landscape of legume genomes, we built a dataset with publicly available genomes of papilionoids and relatives (Supp. Table S3). We then used groups of orthologous genes from the BUSCO dataset to infer a multigene species tree using a coalescent-based approach (Supp. Table S4). The resulting topology was largely consistent with recent phylogenetic studies of Fabaceae (Cardoso et al., 2012; Koenen et al., 2020; Zhao et al., 2021). *N. brasiliensis* is placed within the Dalbergioid clade (Cardoso et al., 2012) and together with its sister species *N. schottii,* it forms a group that is sister to the rest of the clade (Figure 1; Supp. Figure S4).

**Figure 1.**
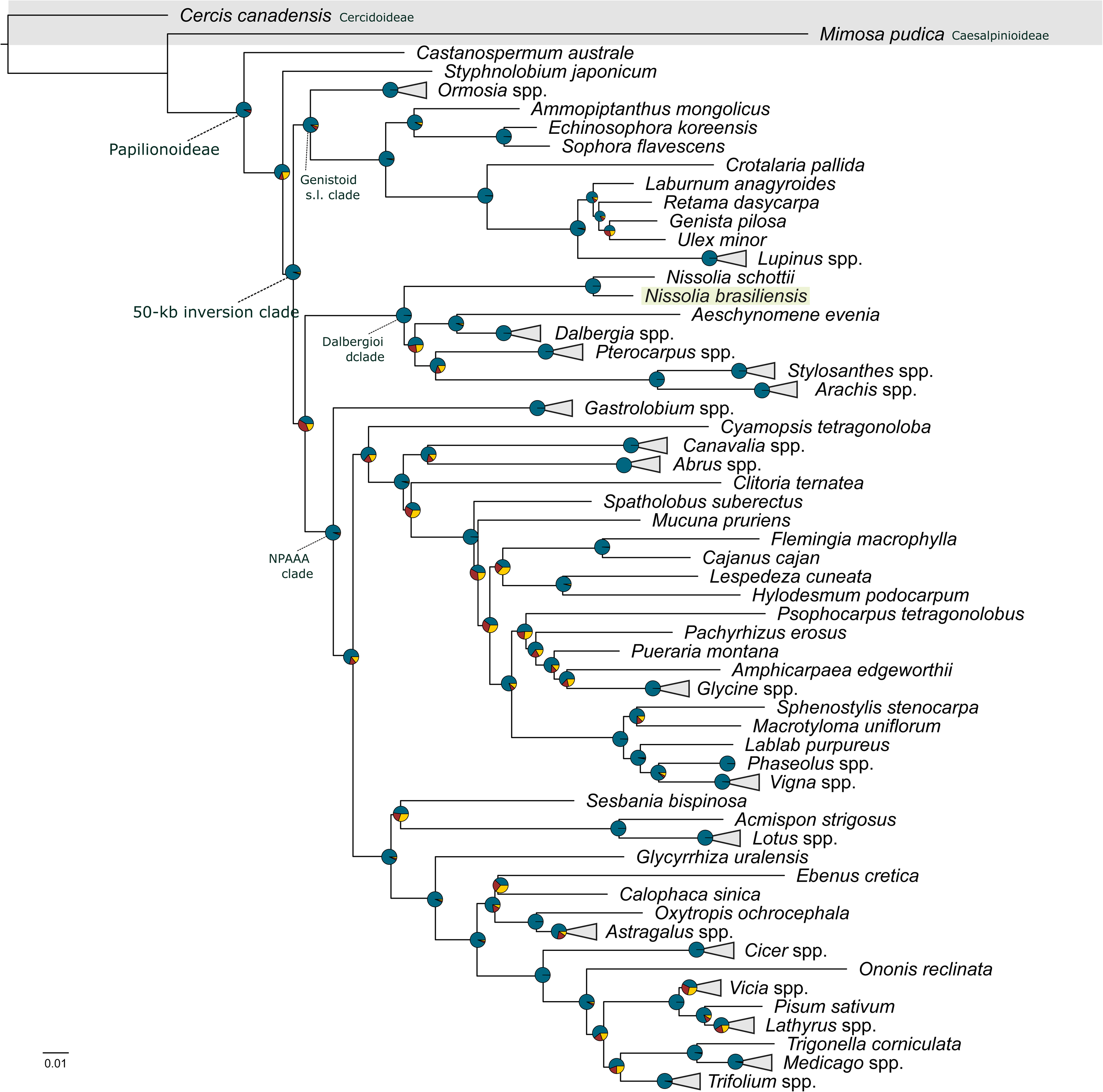
Phylogenetic tree of papilionioid legumes. A multigene species tree was estimated from 1349 genes using a coalescent-based approach. Pie charts on nodes indicate the proportion of quartet trees that support the main topology (dark blue), and the first (red) and second (yellow) alternatives. Local posterior probabilities > 0.9 are omitted from nodes. Genera with multiple representative species are collapsed. Branch lengths are in substitutions per site (Tabatabaee et al., 2023). Taxonomic annotation following (Cardoso et al., 2012). For the full phylogenetic tree, refer to Supp. Figure S4.

The long-read-based *N. brasiliensis* genome presented here covers a part of the legume phylogeny that had been mostly ignored from previous genomic efforts.

### The *Nissolia brasiliensis* genome shows signs of RNS loss

Loss of a trait often correlates with the loss of genes strictly associated with the trait, a process known as co-elimination (Albalat and Cañestro, 2016). In plants, such a co-elimination has been observed in the context of AMS (Delaux et al., 2014; Bravo et al., 2016) and RNS (Griesmann et al., 2018; Van Velzen et al., 2018). Across the NFN clade, the most commonly lost genes following the loss of RNS are the master RNS regulator *Nodule INception* (*NIN*, Schauser et al., 1999) and the infection-related gene *Rhizobium-directed Polar Growth* (*RPG*, Arrighi et al., 2008). Neither *NIN* nor *RPG* were detected in the short read-based assembly of *N. schottii* (Griesmann et al., 2018). Absence of these two genes would further confirm the non-RNS-forming status of *N. brasiliensis*. We conducted a phylogenetic analysis of both genes on a sampling of 25 legume species. *NIN* was clearly identified in all RNS-forming species, but not in the three non-RNS-forming species, including *N. brasiliensis* (Figure 2). Similarly, *RPG* was not detected in any of the three non-RNS forming species. However, *RPG* was also missing from the genomes of the dalbergioids *Arachis ipaiensis*, *Arachis hypogeae*, *Arachis duranensis* and *Aeschynomene evenia*, as previously reported (Quilbé et al., 2021). Micro-synteny analyses between *M. truncatula* and *N. brasiliensis* further supported the absence of both genes in *N. brasiliensis* (Supp. Figure S3B and C). By contrast, and as expected, the genes belonging to the common signaling pathway (*SYMRK*, *CCaMK* and *CYCLOPS*) shared by AMS and RNS are present in *N. brasiliensis.* Genes associated with AMS only (*RAD1*, *STR*, *STR2*) were also detected in *N. brasiliensis,* but not in the non-AMS lupin species, as previously described (Supp. Figure S5 and S6, Delaux et al., 2014; Bravo et al., 2016). The absence of *NIN* from the *N. brasiliensis* long-read-based genome assembly confirms its non-RNS-forming status.

**Figure 2.**
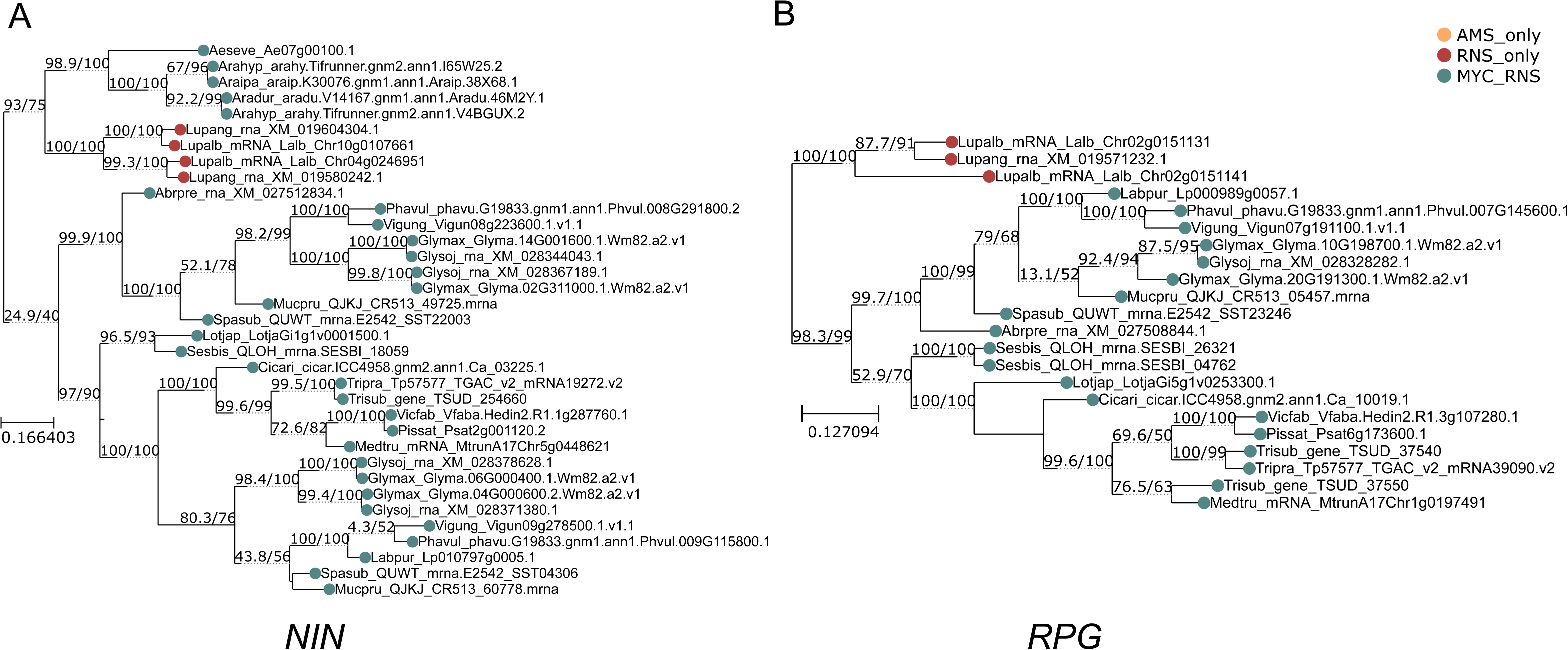
Maximum likelihood phylogenetic trees of *NIN* (A) and *RPG (B)* symbiotic genes. Leaves extremities are colored (yellow, red, grey) according the symbiotic abilities (AMS only, RNS only, AMS and RNS) of each species. Full names of species are available in Supp. Table S4

### Comparative phylogenomics identifies putative RNS genes in Fabaceae

Previous comparative phylogenomic studies included the genomes of RNS and non-RNS forming species within the four orders of the NFN clade: the Fabales, Fagales, Cucurbitales and Rosales. These analyses revealed convergent gene losses associated with the loss of RNS (Griesmann et al., 2018; Van Velzen et al., 2018; Zhang et al., 2024). RNS originated presumably from genetic changes resulting in the rewiring of existing pathways, illustrated by an inferred ancestral RNS transcriptome comprising hundreds of genes (Libourel et al., 2023). Following this initial event, RNS diversified in every lineage giving rise to specific symbiotic programmes (Libourel et al., 2023). The transition from a *Frankia* symbiont to a rhizobia at the base of the legumes, or the convergent evolution of the released symbiosome in Papilionoideae and Mimosoideae legumes are examples of these lineage-specific symbiotic features (Van Velzen et al., 2018; De Faria et al., 2022). To identify genes associated with RNS in legumes, we computed orthogroups for 25 species (Supp. Table S5) and searched for convergent gene losses in the two non-RNS forming lineages: *Castanospermum australe* and the *Nissolia* genus (*N. brasiliensis* and *N. schottii*) and conservation in at least 80% of the nodulating species. This conservation rate (80%) was setup to take into account potential assembly and annotation bias.

A total of 22 genes were retrieved using this filtering, including *NIN* and *RPG*. The other 20 genes covered a large diversity of protein domain annotations (InterPro), ranging from NLR to sugar/inositol transporters and transcription factors (Supp. Table S6). Nine of them were found up-regulated during RNS in at least 50% of the species with available RNA-seq data (Figure 3, Supp. Table S6). Because these 20 genes were not detected in previous phylogenomic analyses conducted at the scale of the NFN clade, their potential symbiotic function might be linked to specificities of the papilionoid legumes, which can be investigated in the future by reverse genetics in RNS-forming legumes.

**Figure 3.**
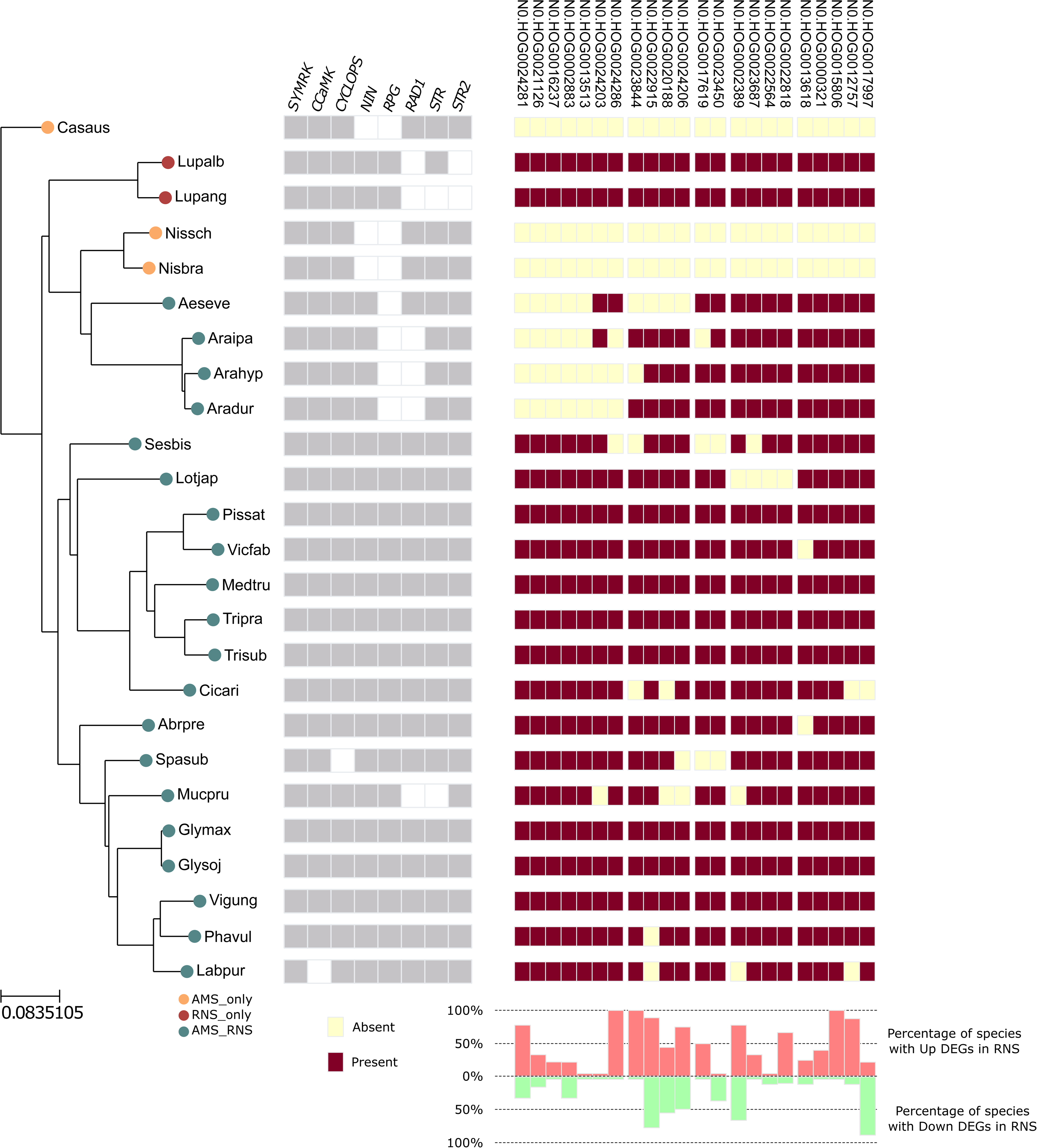
Conservation of symbiotic genes according to symbiotic status in papilonoid species. The tree on the left depicts the papilionoid phylogeny with *Castanospermum australe* as outgroup. The type of symbiosis formed by each species investigated is indicated by a colored dot on the right of each species code name. The heatmap indicates the phylogenetic pattern of presence/absence of genes for each of the 22 investigated orthogroups. The barplots at the bottom indicate the proportion of species that overexpressed (light red) and underexpressed (light green) genes within these orthogroups based on (Libourel et al., 2023) (Supp. Table S6).

### Reduction in symbiotic functions results in convergent gene losses

While the *Nissolia* genus has lost the ability to engage in RNS, the *Lupinus* genus has lost the more ancient AMS (Oba et al., 2001). Although the two lineages lost different types of symbiosis, they both derive from an ancestral state where the two intracellular symbioses, AMS and RNS, co-occurred. How plants, and in particular legumes, deal with two symbioses is poorly understood. Similar to the loss of RNS and AMS, we hypothesised that the transition from a two-symbioses state to a single symbiosis state would lead to the loss of genes specifically involved in the accommodation of two symbionts. Using the previously computed orthogroups (Supp. Table S7) we screened for genes maintained in most legumes but lost in both the *Nissolia* and the *Lupinus* genera while being conserved in at least 80% of the other species. A total of six orthogroups fulfilled these criteria. While all of these genes were lost in *Nissolia*, *Castanospermum* and *Lupinus* (Supp. Table S8), four of them were also lost in *Aeschynomene* and most *Arachis* species, both genera that engage in both RNS and AMS. In contrast with most other Papilionoideae legumes that are infected *via* root-hair infection threads, species from the *Aeschynomene* and *Arachis* genera, as well as *Lupinus*, are infected via an alternative mode of epidermal infection called crack entry (Quilbé et al., 2022). It can be hypothesized that the four genes have been lost in these various lineages following their shifts in symbiotic abilities: *Nissolia* and *Castanospermum* have lost RNS, while *Lupinus*, *Arachis* and *Aeschynomene* have lost the root-hair based infection. Supporting this hypothesis, one of the orthogroup lost in most of these species contains Pectin esterases expressed during RNS in *Medicago* (N0.HOG0024603). Such proteins are known to contribute to cell-wall remodeling during infection and cell-to-cell progression of the infection thread (Su et al., 2023).

Reduction from dual to single symbiotic abilities is not associated to major genomic changes, yet several genes might be linked with a reduction in symbiotic capabilities.

### Stable genetic transformation of *Nissolia brasiliensis*

With its nuclear genome sequenced to a near chromosome-level quality, *N. brasiliensis* now represents a good model to study the loss of RNS. Chimeric root transformation has been previously conducted in the species (Vernié et al., 2025). To further improve the potential of *N. brasiliensis* as a model species, we aimed at developing a stable genetic transformation protocol.

First, we established the different steps of regeneration by somatic embryogenesis without transformation, using different culture media (Table 3). To induce callogenesis, a basic medium (Chabaud et al., 2006) was chosen and four hormone balances were tested (Chabaud et al., 2006; Dan et al., 2006 Supp. Table S9). Three of the tested media allowed calli development on 100% of the explants, with the MR1 medium providing the fastest growth. The sensitivity to Hygromycin was tested during callogenesis on ten explants per condition at the following concentrations: 0mg/L; 5mg/L; 10mg/L; 50mg/L. From 10mg/L, callus formation was no longer observed.

**Table 3.** Regeneration rates of *N. brasiliensis* observed on different media for each stage of regeneration (callogenesis, embryogenesis, rooting). Callogenesis is defined by the number of explants that have produced calli, embryogenesis by the number of calli that have induced shoots and Rooting by the number of shoots that have rooted.

| Callogenesis |  |  |  |  |
| --- | --- | --- | --- | --- |
|  | MR1<br>Auxin: 16µM 2,4-D<br>Cytokinin: 2µM BAP | MR2<br>Auxin: 2,6µM NAA<br>Cytokinin: 2µM BAP | MR3<br>Auxin: 4,5µM 2,4-D<br>Cytokinin: 9,1µM trans-zeatin | MR4<br>Auxin: 0,5µM IAA<br>Cytokinin: 9,1µM trans-zeatin |
| Regeneration rate | 144/144 | 320/320 | 40/40 | 0/40 |

| Embryogenesis |  |  |  |  |
| --- | --- | --- | --- | --- |
|  | ME1 | ME2 | ME3 | ME4 |
| Regeneration rate | 0/126 | 0/126 | 35/126 | 0/126 |

| Rooting |  |  |  |  |  |
| --- | --- | --- | --- | --- | --- |
|  | Mroot1<br>Auxin: 4,9µM IBA | Mroot1+ sucrose<br>Auxin: 4,9µM IBA<br>Sucrose: 20g/L | Mroot2<br>Auxin: 0,9µM IBA | Mroot3<br>No hormones | Mroot4<br>Auxin: 4,9µM IBA<br>+ 0,7µM NAA |
| Regeneration rate | 6/24 | 17/24 | 5/48 | 5/40 | 0/30 |

In a second phase, four embryogenesis media (Chabaud et al., 2006; Mano et al., 2014 Supp. Table S9) were tested for shoot regeneration. Only one of them, the ME3 medium (Mano et al., 2014), was able to induce shoots within six weeks. Finally, the induction of root development was tested using five hormonal conditions (Dan et al., 2006; Mano et al., 2014; Van Velzen et al., 2018 Supp. Table S9). The Mroot1+sucrose medium was proven to be most effective (71% successful root induction). The finalised protocol takes 11 weeks from explant preparation to plant rooting (Figure 4A). During the shooting process, accumulation of a brown pigment in the medium around some regenerating calli was observed and was caused by oxidative stress (Mante and Tepper, 1983). These calli never regenerated shoots and ultimately died, despite being transferred to the ME3 noS medium (adapted from Mante and Tepper, 1983, Supp. Table S9).

**Figure 4.**
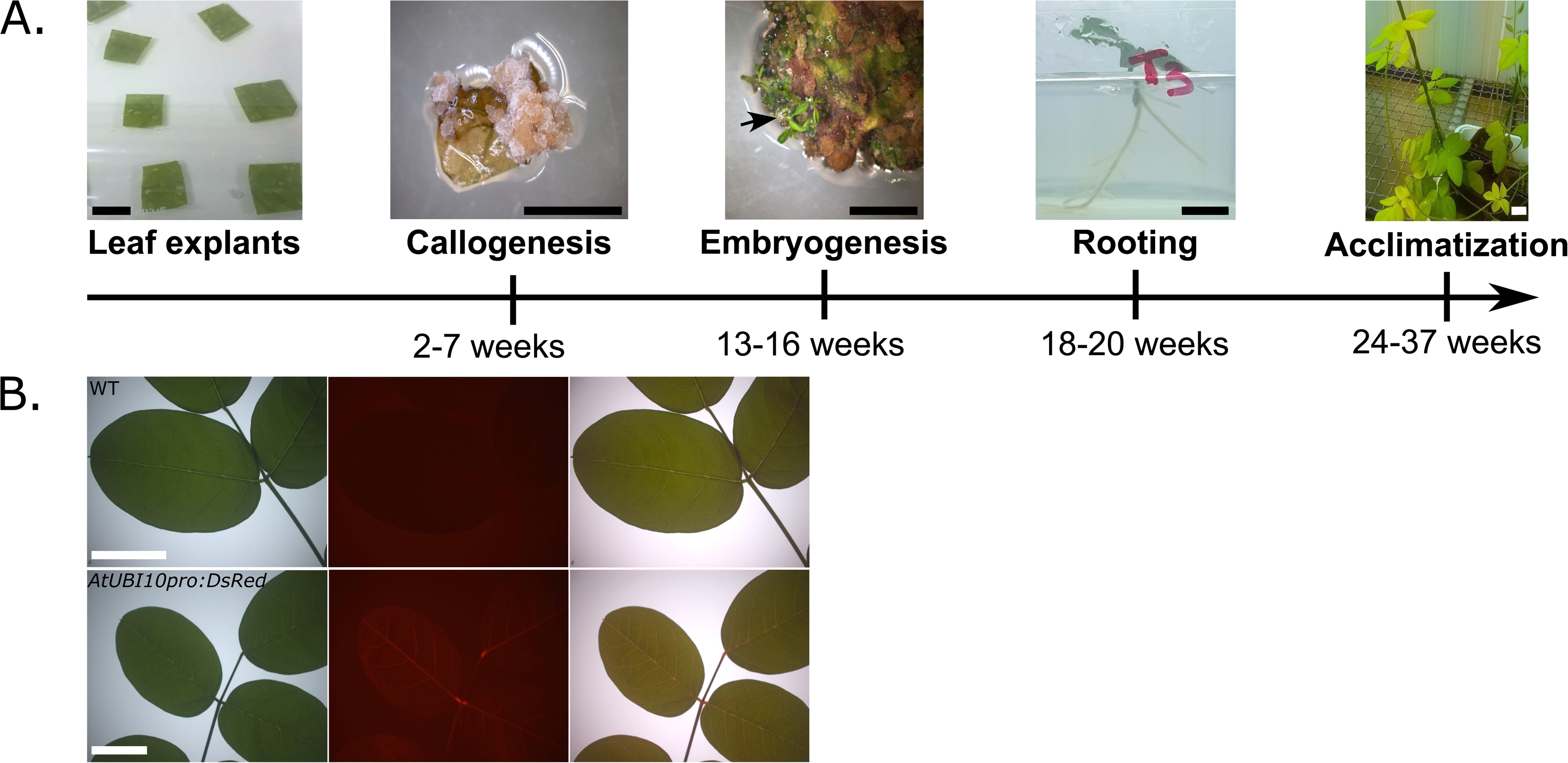
*Nissolia* regeneration protocol. **(A)** Timeline. The protocol starts with leaves explants (a), then callogenesis is induced, with callus visible from 12 days and reaching full size after seven weeks. Calli are transferred to a new medium to induce embryogenesis (c). The first shoots are visible after 42 days, *i.e.* 13 weeks since leaves explants. After 20 days, shoots reach a height of 2 cm, and are transferred to a rooting medium (d). Rooting starts 15 days after transfer and lasts for a further 14 days. (e) Young plantlets are moved on soil with high humidity level to acclimatize, which takes around 30 days and up to 3 months longer to develop into a large plant. Scale bars represent 1 cm (black) or 4 cm (white). **(B)** DsRed fluorescent expression in leaves of a wild-type *N. brasiliensis* plant and a *AtUBI10pro:DsRed*-transformed plant. Bright field, DsRed fluorescence and overlays are shown. Scale bars represent 1 cm.

In our growth conditions, flowers were rare and never produced any seeds. However, vegetative propagation by cuttings was easily achieved (69% survival rate). To do this, young stems 10-15 cm long with terminal buds and axillary buds were cut and transferred to potting soil (see Material and method). After an average of 35 days, roots emerged and the plants could be transferred to larger pots, where they developed into large plants.

Stable genetic transformation has been achieved in a number of legume species (Handberg and Stougaard, 1992; McKently et al., 1995; Olhoft et al., 2003; Chabaud et al., 2006; Mano et al., 2014; Tisseyre et al., 2024), mostly using *Agrobacterium tumefaciens*-mediated delivery of DNA. We therefore used this technique to produce stable transformants of *N. brasiliensis*. As a proof of concept, we created a *N. brasiliensis* line overexpressing DsRed, a fluorescent marker. To do so, explants were incubated with *A. tumefaciens* culture prior to callogenesis conducted on selective media (10 mg/L Hygromycin). Transformed plantlet regenerated and were acclimatised in the greenhouse. The resulting mature plants were genotyped, confirming presence of the DsRED transgene, and DsRed fluorescence was observed in the leaves (Figure 4B).

The stable transformation protocol developed here takes approximately 7-9 months from leaf explants to mature plants to be vegetatively propagated.

## Discussion

Since the sequencing of the first legume genomes – *Medicago truncatula, Glycine max* and *Lotus japonicus* – fifteen years ago, genome assemblies have been generated for more than 100 legume species. The recent transition to long-read-based genome assemblies has revolutionized the field, yet the phylogenetic diversity of the sequenced species remains limited to few, oversampled, legume tribes. For instance, the Trifolieae *Medicago* or *Trifolium* have multiple sequenced species, as does the tribe Phaseoleae with the genera *Vigna*, *Glycine* (including soybean) and *Phaseolus* (including bean). From this initial focus on legume crops and their close relatives, additional clades have been targeted, including the Caesalpinioideae *Mimosa pudica* (Libourel et al., 2023), and the dalbergioids *Aeschynomene aevenia* (Quilbé et al., 2021) and *Arachis hypogaea* (Quilbé et al., 2022). Here, we provide the genome of *N. brasiliensis,* which belongs to a group that is sister to all the other sequenced Dalbergioids (15 species), therefore filling an important gap in the legume genomic landscape. Multiple other papilionoid legume tribes are yet to be investigated at the whole-genome level. Existing large-scale sequencing effort such as 10KP (Cheng et al., 2018; Twyford, 2018) should fill the remaining gaps in the coming decade.

Compared to most other papilionoid legumes, *Nissolia* has lost RNS. Previous sequencing effort had provided two short-read-based genome assemblies for non-RNS forming papilionoids, namely *Castanospermum australe* and *N. schottii* (Griesmann et al., 2018). Although highly fragmentary (N50 = 179 kbp for *N. schottii*), these genomes had already facilitated the identification of two diagnostic genes for the loss of RNS: *NIN* and *RPG* (Griesmann et al., 2018). A denser sampling (125 genome assemblies) of RNS-forming papilionoids, and a much more complete *Nissolia* genome (*N. brasiliensis* N50_18.1 Mbp) allowed us to revisit the initial phylogenomics, focusing on papilionoids. In addition to *NIN and RPG*, another 20 genes were identified. Although it will be interesting to understand the symbiotic function of these genes, using reverse genetics in the model legumes *M. truncatula* or *L. japonicus*, this finding already yields one important conclusion. While the initial gain of RNS before the radiation of the NFN clade resulted in the commitment of at least two genes – *NIN* and *RPG* – to symbiosis, the independent refinement of RNS in the different clades likely resulted in the evolution of clade-specific symbiotic innovations. This had been already exemplified at the functional level by the switch from the ancestral *Frankia* symbiont to rhizobia in legumes, or by the evolution of the symbiosome in (De Faria et al., 2022), and the gene-expression level by the occurrence of species- and clade-specific transcriptomic signatures in response to the symbiont (Libourel et al., 2023). Here, we provide further phylogenomic evidence in legumes. It will be important to explore similar clade-specific evolutionary patterns in the other NFN clades. Rosales, that encompasses a large diversity of RNS-forming and non-RNS-forming species would represent an ideal context for such an analysis.

The engineering of RNS in crops represents one of the most remarkable opportunities in plant synthetic biology. Not only could it boost crop productivity worldwide, it would also strongly reduce the need for nitrogen fertilizers, which have massive ecological impacts (Menegat et al., 2022). Before moving outside the NFN, one particular milestone would be to engineer, or re-evolve, RNS in species belonging to lineages that once had functioning RNS. Such re-evolution would consist in repairing rather than fully engineering, a concept proposed earlier in the context of the Rosales *Trema orientalis*, a close relative of the RNS-forming *Parasponia andersonii* (Behm et al., 2014; Van Velzen et al., 2018). As *N. brasiliensis* is more closely related to the model legumes which have been the main source of knowledge on the genetics of RNS, repairing RNS in that species might be – to date – the shortest path toward engineering RNS. The stable transformation protocol developed here, allowing to generate stable plants in 7-9 months, will allow testing first hypotheses such as the reintroduction of a functional *NIN*.

## Conclusion

Most papilionoid legumes form the nitrogen-fixing root-nodule symbiosis. Here, we have sequenced and assembled a near chromosome-level genome for the non-RNS forming papilionoid legume *N. brasiliensis*. This new genome allowed us to identify 22 candidate genes associated with RNS in legumes. Combining this genome sampling in the legumes with an extended sampling in the other orders of the NFN clade opens the possibility to identify more genes that discriminate between RNS-forming and non-RNS-forming species. Engineering nitrogen fixation in crops species outside the NFN clade is a major endeavour that would transform agriculture (Bailey-Serres et al., 2019; Jhu and Oldroyd, 2023). The development of the first stable transformation protocol for a non-RNS-forming legume offers an interesting possibility in that direction: the re-evolution of RNS by introducing lost genes in this close relative of the model legumes *Medicago truncatula* and *Lotus japonicus*.

## Acknowledgements

The authors would like to thank Renan Destrade, Wandrille Delaporte, Anaïs Van Den Berghe and Xavier Bossier for the great maintenance of plants in all circumstances. Also Corinne Lefort for her help with maintaining the transgenic material and finally Annabelle Dupas for helpful discussions and technical assistance. We are grateful to the genotoul bioinformatics platform Toulouse Occitanie (Bioinfo Genotoul, https://doi.org/10.15454/1.5572369328961167E12) for providing computing resources. This study was supported by the Fédération de Recherche Agrobiosciences, Interactions et Biodiversité, the “Laboratoires d’Excellence (LABEX)” TULIP (ANR-10-LABX-41)”. This work was supported by the project Engineering Nitrogen Symbiosis for Africa (ENSA) currently funded through a grant to the University of Cambridge by the Bill & Melinda Gates Foundation (OPP1172165) and the UK Foreign, Commonwealth and Development Office as Engineering Nitrogen Symbiosis for Africa (OPP1172165). MEB is supported by the Horizon Europe programme MSCA-PF (grant 101105838 ‘SYMBIOLOSS’).

## Data availability

Genome assembly and RNASeq reads have been deposited in the ENA-EMBL platform under the accession number: PRJEB90298. Raw genomic reads have been deposited in the NCBI under the accession number: PRJNA736193.

A genome browser of *N. brasiliensis* is accessible at: https://bbric-pipelines.toulouse.inra.fr/myGenomeBrowser?browse=1&portalname=Nissolia_brasiliensis&owner=jean.keller@cnrs.fr&key=UKsIw0Sf

## Material and method

### Biological material

Seeds of *N. brasiliensis* (accession number: 13435) were obtained from the International Center for Tropical Agriculture (CIAT, Cali, Colombia). For germination, seeds of *N. brasiliensis* and *M. truncatula* A17 were scarified with pure sulfuric acid for 7min. After one min in pure bleach (9.6% chl) and five washes, seeds were placed on inverted agar plates (0,8%) for three days in the dark at 8°C and then for five days at 25°C/ photoperiod 16h-8h/ 65% humidity. For *Lotus japonicus* Gifu, seeds were scratched with sandpaper until whitened. After fifteen min in diluted bleach (1:15 v/v) and 3 washes, seeds were left to soak for thirty min then placed on inverted 0.8% agar plates for 3 days in the dark at 21°C/ photoperiod 16h-8h/ 65% humidity.

### Cutting protocol

Young stems (10-15 cm long) with terminal buds and axillary buds were cut and transferred in zeolite substrate (50% fraction 1.0 to 2.5 mm, 50% fraction 0.5 to 1.0- mm, Symbiom) at 22°C, 70% humidity, photoperiod 16h/8h. Once roots emerged and the plants could be transferred to larger pots, where they developed into large plants.

### Nodulation assay

Rhizobia strains (*S. meliloti* RCR2011 pXLGD4 (GMI6526), *R. tropici* CIAT 899 phC60*, S. fredii* NGR234 phC60*, IRBG74* pSKDSRED*, M.loti R7A* pSKDSRED) (Montiel et al., 2021) were grown for two days at 28°C in liquid tryptone yeast medium supplemented with 6mM calcium chloride, rinsed with water and diluted to OD_600_=0.02. Each pot was inoculated with 10mL of bacterial suspension and plants were watered with a Fahreus solution (Catoira et al., 2000). IRBG74 was inoculated on germinated seedlings of *N.brasiliensis*, whereas other rhizobial strains were inoculated on cuttings of *N. brasiliensis*. Plants were grown on sterile zeolite substrate (50% fraction 1.0 to 2.5 mm, 50% fraction 0.5 to 1.0- mm, Symbiom) under the conditions 22°C, 70% humidity, photoperiod 16h/8h and nodules were observed thirty to forty days post inoculation. Seedlings of *M. truncatula* and *L. japonicus* were used as control plants.

### Arbuscular mycorrhizal symbiosis assay

Germinated seedlings were inoculated with 500 spores of Ri DAOM197198 (Agronutrition AP-2007, Toulouse, France) by pot and cultivated during 7 weeks at 20°C/ photoperiod 16h-8h/ 70% humidity in sterilized zeolite substrate (50% fraction 1.0 to 2.5 mm, 50% fraction 0.5 to 1.0- mm, Symbiom). Twice per week, plants were watered with a solution of Long Ashton Low P (Hewitt, 1953).

Mycorrhization root colonization was observed after ink staining. Root systems were incubated for eight min at 95°C in 10% (w/v) KOH, rinsed thoroughly with water, incubated for eight min at 95°C in an ink staining solution (5% Sheaffer ink, 5% acetic acid) and rinsed thoroughly with water again.

### High-molecular weight DNA extraction

Plants were grown on mixture soil (mix of sphagnum peat/ clay/ pearlstone provided by Proven® Substrate) at 20°C/ photoperiod 16h-8h/ 70% humidity and watered once time with fertilizer per week.

DNA was isolated from dark treated young leaves using QIAGEN Genomic-tips 100/G kit (Cat No./ID: 10243) following the tissue protocol extraction. Briefly, 1g of young leaf material were frozen and ground in liquid nitrogen with mortar and pestle. After 3h of lysis at 50°C and one centrifugation step, the DNA was immobilized on the column. After several washing steps, DNA is eluted from the column, then desalted and concentrated by alcohol precipitation. The DNA is resuspended in EB buffer. DNA quality and quantity were assessed respectively using the Nanodrop-one spectrophotometer (Thermo Scientific) and the Qbit 3 Fluorometer using the Qbit dsDNA BR assay (Invitrogen). The size of the DNA was assessed using the FemtoPulse system (Agilent, Santa Clara, CA, USA).

### Library preparation

An HIFI SMRTbell® library was constructed using the SMRTbell® Template Prep kit 2.0 (Pacific Biosciences, Menlo Park, CA, USA) according to PacBio recommendations (PN 101-853-100, version 05). Briefly, HMW DNA was sheared by using Megaruptor 2 system (Diagenode, Liège Science Park, Belgium) to obtain a 20Kb average size. Following an enzymatic treatment on 10µg of sheared DNA sample for removing single-strand overhangs and DNA damage repair, ligation with overhang adapters to both ends of the targeted double-stranded DNA (dsDNA) molecule was performed to create a closed, single-stranded circular DNA. A nuclease treatment was performed by using SMRTbell® Enzyme Clean-up kit 1.0 (Pacific Biosciences, Menlo Park, CA, USA). A size-selection with Blue-Pippin system (Sage Science, Beverly, MA, USA) to remove fragments less than 10Kb was done on purified sample with 1X AMPure PB beads (Pacific Biosciences, Menlo Park, CA, USA). The size and concentration of the final library were assessed using the FemtoPulse system (Agilent, Santa Clara, CA, USA) and the Qubit Fluorometer and Qubit dsDNA HS reagents Assay kit (Thermo Fisher Scientific, Waltham, MA, USA), respectively.

Sequencing primer v2 and Sequel® II DNA Polymerase 2.0 were annealed and bound, respectively to the SMRTbell library. The library was loaded on 2 SMRTcells 8M at an on-plate concentration of 70pM using a diffusion loading. Sequencing was performed on the Sequel® II system with Sequel® II Sequencing kit 2.0, a run movie time of 30 hours with 120 min pre-extension step and Software version 9.0 PacBio) by Gentyane Genomic Platform (INRAE-Clermont-Ferrand, France).

### PacBio Hi-Fi sequencing and assembly

The data were assembled using HiFiasm assembler (v0.13; (Cheng et al., 2021). Hifiasm is able to produce a primary assembly and alternative assembly (incomplete alternative assembly consisting of haplotigs under heterozygous regions. All the metrics correspond to the primary assembly used for the next analysis. We obtained 1,647,178 corrected reads (N50 = 18.1Mbp; Genome Coverage = 33X) assembled in 654 contigs (N50 = 37,5Mbp; L50 = 8 contigs; GC content = 37.26%). A BUSCO analysis was performed with the viridiplantae database in order to evaluate the quality of the assembly produced (Simão et al., 2015). We obtained a 98.6% complete BUSCO score on the assembly.

### Preparation of ultra-high molecular weight (uHMW) DNA for Bionano optical mapping

uHMW DNA were purified from 1g of fresh dark treated very young leaves according to the Bionano Prep Plant Tissue DNA Isolation Base Protocol (30068 - Bionano Genomics) with the following specifications and modifications. Briefly, the leaves were fixed in fixing buffer containing formaldehyde. After 3 washes, leaves were cut in 2mm pieces and disrupt with rotor stator in homogenization buffer containing spermine, spermidine and beta-mercaptoethanol. Nuclei were washed, purified using a density gradient and then embedded in agarose plugs. After overnight proteinase K digestion (Qiagen) in the presence of Lysis Buffer and one hour treatment with RNAse A (Qiagen), plugs were washed and solubilized with 2 µL of 0.5 U/µL AGARase enzyme (ThermoFisher Scientific). A dialysis step was performed in TE Buffer (ThermoFisher Scientific) to purify DNA from any residues. The DNA samples were quantified by using the Qubit dsDNA BR Assay (Invitrogen). The presence of mega base size DNA was visualized by pulsed field gel electrophoresis (PFGE).

Labeling and staining of the uHMW DNA were performed according to the Bionano Prep Direct Label and Stain (DLS) protocol (30206 - Bionano Genomics). Briefly, labeling was performed by incubating 750 ng genomic DNA with DLE-1 enzyme (recognizing the site CTTAAG) in the presence of DL-Green dye. Following proteinase K digestion and DL-Green cleanup by membrane adsorption, the DNA backbone was stained by mixing the labeled DNA with DNA Stain solution and incubated overnight at room temperature. The DLS DNA concentration was assessed with the Qubit dsDNA HS Assay (Invitrogen).

### Data collection, optical mapping construction and hybrid assembly

Labeled and stained DNA was loaded on the Saphyr chip. Loading of the chip and running of the Bionano Genomics Saphyr System were performed according to the Saphyr System User Guide (30247 - Bionano Genomics). Data processing was performed using the Bionano Acess and Solve v.3.6 software (https://bionano.com/software-and-data-analysis-support/).

A total of 437 Gb filtered data were generated (molecules with a size larger than 150kb) corresponding to 437X coverage of the estimated size of the genome. The assembled data represent 137 maps with a N50 of 28Mbp and a total length of 915 Mbp.

Finally, a hybrid scaffolding was performed between the PacBio assembly (HiFiasm) and the optical genome maps using the Bionano Hybrid Scaffold pipeline with default parameters. We obtained 37 hybrid scaffolds with a N50 of 35 Mbp and a total length of 752 Mbp (maximum scaffold size = 90Mbp).

### Genome size and chromosome number estimations

#### Genome size estimation

Approximately four mg of fresh leaves from *N. Brasiliensis* and the reference model plant *Z. mays* (2C DNA = 5.55 pg, (Zonneveld, 2019)) were harvested and transferred to a petri dish. Estimation of genome size for each species was obtained as described by (Boutte et al., 2020). Three measures of genome size were made. From each, the mean ratio of DNA content was calculated. Genome sizes were converted from picograms (pg) to Megabases (Mb) using 1 pg = 978 Mbp (Doležel et al., 2003).

#### Chromosome counting

The chromosome number of *N. brasiliensis* were determined from mitotic chromosomes observed on metaphasic cells isolated from root tips. Roots tips of 0.5 - 1.5 cm length were treated in the dark with 0,04% 8-hydroxiquinoline for 2h at 4°C followed by 2h at room temperature to accumulate metaphases. They were then fixed in 3:1 ethanol-glacial acetic acid for 12 hours at 4°C and stored in ethanol 70 % at -20 °C until use. After being washed in distilled water for 10 min, then treated 15 min by a 0.01 M citric acid-sodium citrate pH 4.5 buffer, root tips were incubated at 37°C for 30 min in a enzymatic mixture (5% Onozuka R-10 cellulase (Sigma), 1% Y23 pectolyase (Sigma)). The enzymatic solution was removed and the digested root tips were then carefully washed with distilled water for 30 min. One root tip was transferred to a slide and macerated with a drop of 3:1 fixation solution. Dried slides were then stained by a drop of 4’,6-diamidino-2-phenylindole (DAPI). Cells were viewed with a ORCA-Flash4 (Hamamatsu, Japan) on an Axioplan 2 microscope (Zeiss, Oberkochen, Germany) and analysed using Zen software (Carl Zeiss, Germany).

#### RNA-sequencing

Plants were obtained by cuttings and grown on sterilized zeolite substrate (50% fraction 1.0 to 2.5 mm, 50% fraction 0.5 to 1.0- mm, Symbiom) at 22°C, 50% humidity, photoperiod 16h/8h, watered twice per week with a solution of Long Ashton Low P (Hewitt, 1953)). RNA extraction and DNase treatment were performed respectively using E.Z.N.A. RNA extraction kit (Omega-Biotek) and TURBO DNA-free kit (Invitrogen) according to manufacturers’ instructions on leaves, stems, uninoculated roots and seven weeks old mycorrhized roots. Quality of RNAs was assessed using the Agilent 2100 Bioanalyzer system. RNA sequencing was performed by the Eurofins genomics facility. Polyadenylated mRNA and RNA-seq libraries were prepared according to Illumina’s protocols using the Illumina TruSeq Stranded mRNA sample prep kit to analyse mRNA. RNA-seq experiments were performed on an Illumina NovaSeq 6000 using a paired-end read length of 2 × 150 bp with the Illumina NovaSeq 6000 sequencing kits.

### Genome annotation

Genome of *N. brasiliensis* has been first soft-masked using the RepeatMasker tools suite. Repeats were predicted using RepeatModeler v2.0.6 (Flynn et al., 2020) with the rmblast 2.14.1+ as search engine and the LTR structural analysis enabled. Predicted repeats were then used to soft-mask the genome using RepeatMasker v4.1.7-p1 (http://www.repeatmasker.org) and the same search engine as in RepeatModeler. To optimize the gene predictions, RNA-Seq data have been produced as described above. For each sample, reads were first trimmed using TrimGalore v0.6.10 (Krueger et al., 2023) with reads length and quality threshold set up to 50bp and 30 respectively. Processed reads were then aligned against the *N. brasiliensis* genome using HISAT2 v2.2.1 with the following paramters: ‘--sensitive --max-intronlen 5000 --no-mixed --no-discordant --dt’. Then, each mapping file were converted into BAM files and processed to group reads by name, fill mate coordinates and then sorted by position before tagging duplicated reads using the view, collate, fiwmate, sort and markdup options from samtools v1.19 (Danecek et al., 2021). One sample was discarded after the mapping due to an alignment rate lower than 10%. Finally, all mappings for each sample were concatenated and treated as the individual ones described above using samtools.

In parallel, a reference protein database was constructed to complement RNA-Seq hints during annotation process. To this aim, the Viridiplantae OrthoDB11 was concatenated with proteomes of 10 papilionoids covering the different phylogenetic clades (see SuppTable X). Both RNA-Seq and protein data described here were provided to BRAKER3.0.7 (Gabriel et al., 2024) along with the sofmasked genome and ‘Fabales’ as BUSCO lineage option to optimize annotation. Predictions from BRAKER were then post-processed to, first, fix overlapping genes and then keep only the longest isoform per gene using respectively agat_sp_fix_overlaping_genes.pl and agat_sp_keep_longest_isoform.pl scripts from the AGAT v1.4.0 (Jacques Dainat et al., 2024). A public genome browser for the genome and its annotation has been made available using the myGenomeBrowser tool (10.1093/bioinformatics/btw800). Finally, protein-coding genes were functionally annotated using InterProScan5 v64.96.0 (Blum et al., 2025) with the ‘-iprlookup -goterms’ parameters enabled. All the synteny analyses presented here have been conducted using the JCVI v1.5.4 (Tang et al., 2024) toolkit, the protein-coding genes and limited to scaffolds contained within the L90 metric of their respective genomes.

### Papilionoid species tree reconstruction

To place *N. brasiliensis* in an evolutionary context, we inferred a phylogenetic tree of papilionoid legumes. Genomes of all papilionoid species available on NCBI (last accessed: May 2025) were retrieved and complemented with genomes from other sources (see Supp.Table S3 and Supp. TableS4), and only a single assembly per species was retained. To obtain shared phylogenetic markers across species, we used BUSCO v5.8.3 (Manni et al., 2021) to search for orthologous protein sequences against the ODB12 Fabales database (7843 markers). Here, two species were removed due to low gene completion rate (below 20%; *Trifolium medium* - GCA_003490085.1 and *Dipterix alata* - GCA_012978445.1), resulting in a set of 119 species. We then extracted all homologs (including paralogs) from all genomes for each marker. To reduce missing data, only markers that were present in at least 70% of the species were retained. We then enriched the data set with single-copy markers, to reduce the impact of divergent paralogs on the phylogenetic analysis. For that, we retained only markers that were present as single-copy genes in at least 80% of the species. Sequences were then aligned using MAFFT v705 (Katoh and Standley, 2013) with default parameters, and alignments were trimmed using trimAl v1.4.1 (Capella-Gutiérrez et al., 2009) to remove columns with more than 50% missing data (-gt 0.5). To reduce the impact of uninformative markers, we removed those with alignment length < 500 bp after trimming. A maximum likelihood phylogenetic tree was then estimated for each of the remaining 1349 markers using IQ-Tree v2.2.2.6 (Minh et al., 2020) with the standard substitution model selection. To remove potential spurious homologs and pseudogenes, we used TreeShrink v1.3.9 (Mai and Mirarab, 2018) to detect abnormally long branches with a false positive rate of 0.05, and these were then excluded from the trees. Finally, a multigene coalescent species tree was inferred on the filtered gene trees using Astral-Pro3 v1.19.3.5 (Zhang and Mirarab, 2022; Tabatabaee et al., 2023).

### Search for symbiotic genes

Key symiotic genes involved in both arbuscular mycorrhization and nodulation (*SYMRK, CCaMK CYCLOPS*) as well as specific to AMS (*STR, STR2, RAD1*) and to RNS (*NIN, RPG*) were searched in *N. brasiliensis*. To this aim, reference protein from the model legume *M. truncatula* were searched against a database composed of proteomes from representative papilionoid species (Supp. Table S3) using the BLASTp v2.16.0+ algorithm (Camacho et al., 2009) with an e-value threshold of 1^e-03^. Homologous sequences were then aligned using MAFFT v7.526 (Katoh and Standley, 2013) and alignments trimmed with trimAl v1.5.rev0 (Capella-Gutiérrez et al., 2009) to remove positions containing more than 60% of gaps. Trimmed alignments were provided to FastTree v2.1.11 (Price et al., 2010) to build a first phylogenetic tree. From these trees, orthologous clade for each query were extracted and a phylogenetic tree rebuilt. To this, orthologous protein were first aligned using PRANK v250331 (available at https://github.com/ariloytynoja/prank-msa) with a maximum of 10 iterations before trimming alignment as described previously. Phylogenetic trees were finally reconstructed using IQ-TREE2 v2.2.2.6 (Minh et al., 2020) with the identification of the best-fitting evolutionary model estimated with ModelFinder (Kalyaanamoorthy et al., 2017) and the Bayesian Information Criteria. Branch supports were tested using 10,000 replicates of both sh-aLRT (Guindon et al., 2010) and UltraFast Bootstraps (Hoang et al., 2018). Trees were vizualised and processed using both the iTOL v7 online platform (Letunic and Bork, 2024) and the python package ete4 v4.3.0 (Huerta-Cepas et al., 2016).

### Comparative phylogenomics and phylotranscriptomics

To investigate the potential gene losses associated with the loss of the RNS in *Nissolia spp.* and *Castanospermum australe*, we reconstructed the orthogroups using OrthoFinder v2.5.5 (Emms and Kelly, 2019) with the ‘-diamond_ultra_sens -M ms’ option to maximize the accuracy of the analysis. To run OrthoFinder, a proteomes database of 25 papilionoid species representing the different phylogenetic clades was used (SuppTable SX). Orthogroups were then filtered to retain only the ones where both *Nissolia* species and *C. australe* were absent and containing at least 80% of the other papilionoid species. The same filter was applied a second time with the additional constraint that the two lupin species (RNS but no AMS) had to be also absent. To further characterize, particularly at the transcriptomic level, these orthogroups, we referred to Libourel et al., 2023, to identify the orthogroups containing the *M. truncatula* genes (Supp. Table S6). We then calculated the percentage of species that overexpressed and underexpressed genes within these orthogroups (Supp. Table S6). The percentages (Perc_NodUp or Perc_NodDown) are calculated by taking the number of species among the 9 that possess an Up and Down, respectively, deregulated gene in the orthogroup divided by the number of species among the 9 that contain a gene in the orthogroup.

### Transformation protocol

#### Cloning

The Golden Gate modular cloning system (Engler et al., 2014; Patron et al., 2015) was used to prepare the plasmids as described by (Rich et al., 2021). DsRed module (EC15073) was clone under *Arabidopsis thaliana Ubiquitin* promoter (EC15062) with a 35S Terminator (EC41414) in EC47811 to create a Level1 plasmid. This Level1 plasmid was associated with EC15030 and EC41744 in EC50507 to create a plasmid overexpressing DsREd and Hygromycin (See (Vernié et al., 2025) for plasmids sequences).

#### Preparation of Agrobacterium culture

*Agrobacterium tumefaciens* (GV3101) containing a construct overexpressing DsRED and Hygromycin resistance gene was inoculated in two mL of liquid LB with selective antibiotics, 48h before transformation. This pre-culture was used to inoculate thirty mL of liquid LB with selective antibiotics, 24 hours before transformation. These liquid cultures were incubated at 28° C with rotary shaking at 180 rpm. *Agrobacterium* culture were then centrifuged at 3000rpm for fifteen min. Pellet was resuspended in liquid MR1-AS medium (Chabaud et al., 2006) to an OD_600_=0.6.

#### Plant material preparation

Young leaves of *N. brasiliensis* grown at 24°C/ photoperiod 16h-8h/ 70% humidity were collected. Leaves were washed with a water/soap solution, then sterilized for three min with a 6% bleach solution and rinsed five times with sterile water. Leaves were then cut to form explants of 1 to 2 square cm.

#### Co-culture and callogenesis

Explants were transferred to a Petri dish containing the *Agrobacterium* suspension and vacuum was applied for thirty min. Explants were then gently shacked for 2h and transferred to MR1-AS solid medium (Supp. Table S9) for 48 h and then to MR1 H10 AUG800 medium (Chabaud et al., 2006), Supp. Table S9) to induce callogenesis and select the transformed explants. Co-culture and callogenesis were performed in the dark, at 24°C and 70% humidity.

#### Whole plant regeneration

Once calli covered the explant, they were transferred on ME3 H10 (Mano et al., 2014) Supp. Table S9) to induce shooting (media was changed monthly). Browning calli were transfered on ME3 H10 noS (Mante and Tepper, 1983; Mano et al., 2014) Supp. Table S9). After shoots apparition, shoot 2cm high were cut at their basis and transferred on Mroot1+sucrose (Dan et al., 2006), supp. Table S9) for rooting. Shooting and rooting were performed at 25°C/ photoperiod 16h-8h/ 65% humidity.

#### Genotyping

DNA extraction was performed following the extraction process of Kapa3G Plant PCR Kit (KapaBiosystems). PCR amplification of DsRed was performed with 2 specific primers (forward 5‘AGGGTTTCATAGATATCATCCG 3’ ; reverse 5‘GTCGATCGACTCTAGCTAGA 3’) and with the following cycle : 95°C for 3min, then 35 cycles of 95°C for 20s, 56°C for 15s, and 72°C for 30sec, with a final extension reaction at 72°C for 1min.

#### DsRed observation

Excitation wavelength used was between 513-556 and emitted wavelength 570-613nm. Images were acquired with a Zeiss Axiozoom V16 microscope.

## Supplementary figures

**Supp. Fig. S1. Nissolia brasiliensis root myccorrhized by Rhizophagus irregularis.**

*Nissolia* roots were inoculated for 7 weeks. *R. irregularis* is visible in dark blue after ink staining. Differents fungal structures were observed: spore (s), hyphae (h), vesicle (v) and arbuscules (a).

**Supp. Fig. S2. Chromosome counting results for three independent replicates of *N. brasiliensis* mitotic root tip.** Each picture corresponds to a different individual (Rep. 1,2,3).

**Supp. Fig. S3.** Synteny analysis between *M.truncatula* and *N.brasiliensis* based on the scaffolds included within the L90 limit. A. Whole genome synteny analysis. B. and C. Synteny analysis of MtNIN and MtRPG regions respectively

**Supp. Fig. S4. Phylogenetic tree of papilionoid legumes.** Multigene species tree estimated from 1349 genes using a coalescent-based approach. Pie charts on nodes indicate the proportion of quartet trees that support the main topology (dark blue), and the first (red) and second (yellow) alternatives. Local posterior probabilities are shown next to nodes. Branch lengths are in substitutions per site (Tabatabaee et al. 2023) (). Taxonomic annotation following (Cardoso et al. 2012) ().

**Supp. Fig. S5. Maximum likelihood phylogenetic trees of symbiotic genes involved in both AMS and RNS (SYMRK, CYLCOPS, CCaMK) and AMS only (STR, STR2, RAD1).** Leaves extremities are colored (yellow, red, grey) according the symbiotic abilities (AMS only, RNS only, AMS and RNS) of each species. Full names of species are available in Supp. Table S4

**Supp. Fig S6. Estimated species tree from OrthoFinder analysis with presence or absence of symbiotic genes** indicated with grey (ortholog present) or empty (ortholog absent) squares. Leaves extremities are colored (yellow, red, grey) according the symbiotic abilities (AMS only, RNS only, AMS and RNS) of each species. Full names of species are available in Supp. Table S4

## Supplementary tables

**Supp. Table S1. Number of plants showing infected nodules in response to five *Rhizobia* species.** IRBG74 inoculation was carried out on germinated *N. brasiliensis* seeds, whereas other *Rhizobia* species were inoculated onto *N. brasiliensis* cuttings. NT=Not Treated. *Medicago truncatula* and *Lotus japonicus* were used as positive controls for nodulation.

**Supp. Table S2.** Genome size estimation on three independent individuals of *N. brasiliensis*. *Z. mays* was used as a reference to estimate the *N. brasiliensis* genome size.

**Supp. Table S3.** List of species obtained from the NCBI to build the species tree. For each species, the BUSCO score has been calculated using BUSCO V5.8.3 and the Fabales ODB12 database (n=7843).

**Supp. Table S4.** List of species included in the OrthoFinder and species tree reconstruction analyses. For each species, the BUSCO score has been calculated using BUSCO V5.8.3 and the Fabales ODB12 database (n=7843).

**Supp. Table S5. Orthogroups absent in the non-nodulating species (*N. schottii, N. brasiliensis* and *C. australe*) while being maintained in at least 80% of RNS-forming species.**

**Supp. Table S6.** Orthogroups from (Libourel et al., 2023) containing the *M. truncatula* genes from Supp. Table S5 with the proportion of species that overexpressed and underexpressed genes within these orthogroups in nodulation condition. The percentage (Perc_NodUp and Perc_NodDown) are calculated by taking the number of species among the 9 that possess an Up and Down, respectively, deregulated gene in the orthogroup divided by the number of species among the 9 that contain a gene in the orthogroup.

**Supp. Table S7. Phylogenetic hierarchical orthogroups computed with OrthoFinder.**

**Supp. Table S8. Orthogroups absent in the non-nodulating species (*N. schottii*, *N. brasiliensis* and *C. australe*) and the two *Lupinus* species while being maintained in at least 80% of other RNS-forming species.**

**Supp. Table S9. Composition of the tissue culture media used for *Nissolia brasiliensis* transformation and regeneration protocols.**

## Notes

### Competing Interest Statement

The authors have declared no competing interest.

