## Supplementary material for "*Nissolia brasiliensis* as a non-nodulating model legume": Figure S1

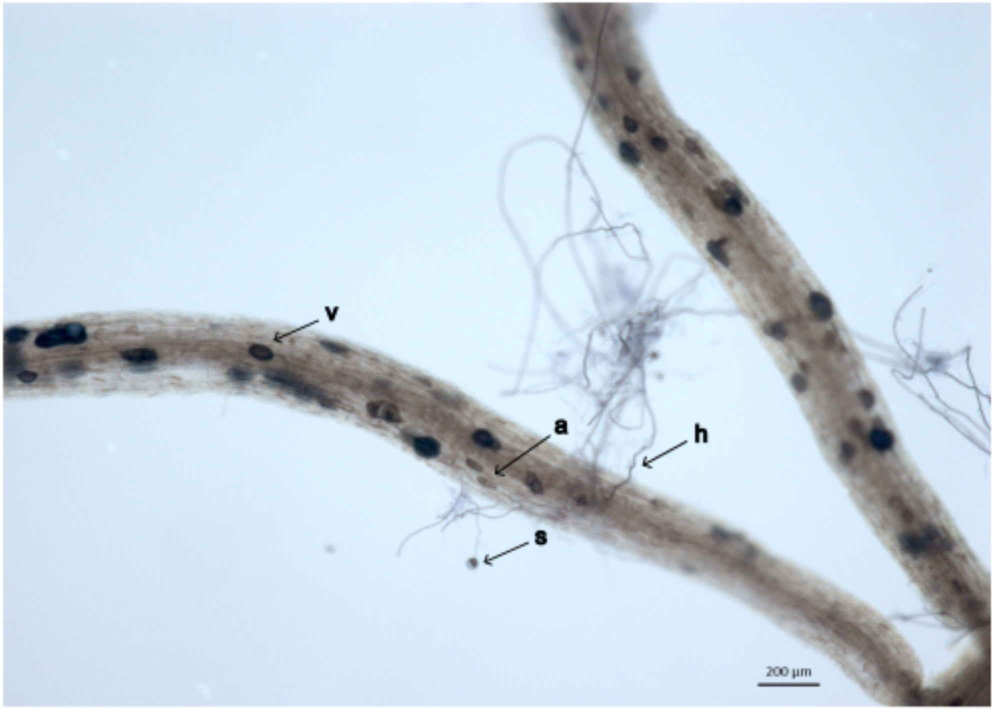


**Supplementary Figure S1.** ***Nissolia brasiliensis*** **root myccorrhized by** ***Rhizophagus irregularis.***

*Nissolia* roots were inoculated for 7 weeks. *R. irregularis* is visible in dark blue after ink staining. Differents fungal structures were observed: spore (s), hyphae (h), vesicle (v) and arbuscules (a).
