## Supplementary material for "*Nissolia brasiliensis* as a non-nodulating model legume": Figure S2

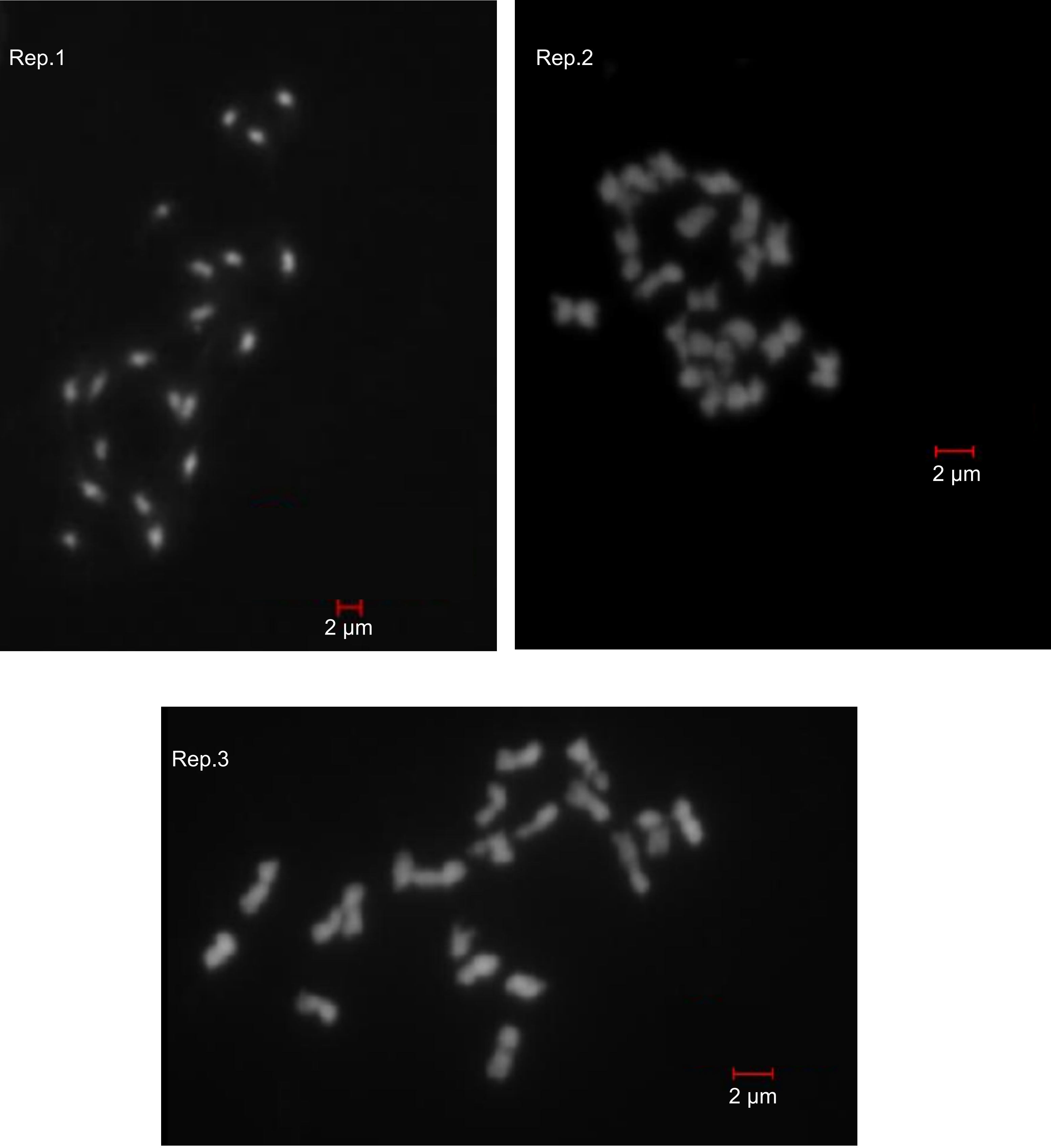


**Supplementary Figure S2.** Chromosome counting results for three independent replicates of *N. brasiliensis* mitotic root tip. Each picture corresponds to a different individual (Rep.1,2,3).
