## Supplementary material for "*Nissolia brasiliensis* as a non-nodulating model legume": Figure S3

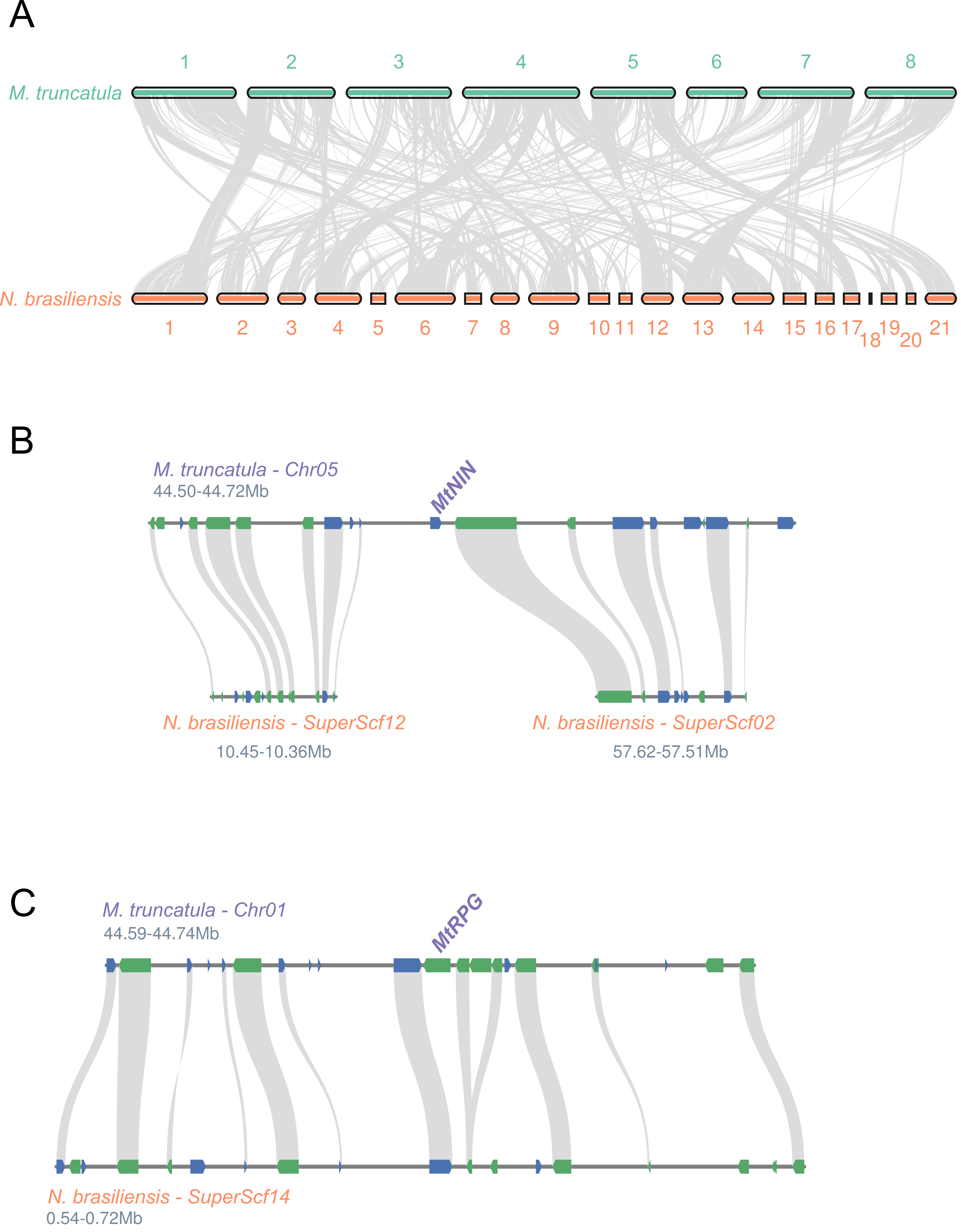


**Supplementary Figure 3. Synteny analysis between *M.truncatula* and *N.brasiliensis* based on the scaffolds included within the L90 limit. A.** Whole genome synteny analysis. **B.** and **C.** Synteny analysis of *MtNIN* and *MtRPG* regions respectively
