## Supplementary figures and images for "*Nissolia brasiliensis* as a non-nodulating model legume"

### Figure S4

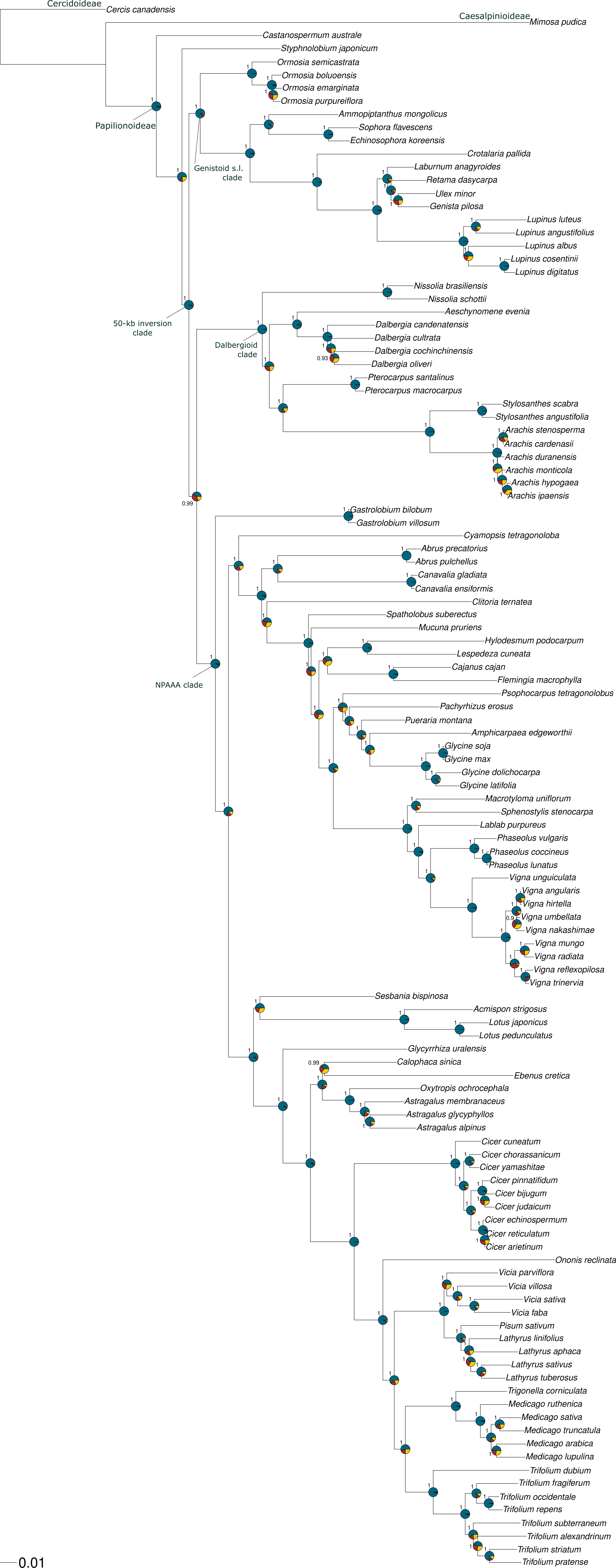
