## Supplementary material for "*Nissolia brasiliensis* as a non-nodulating model legume": Figure S5

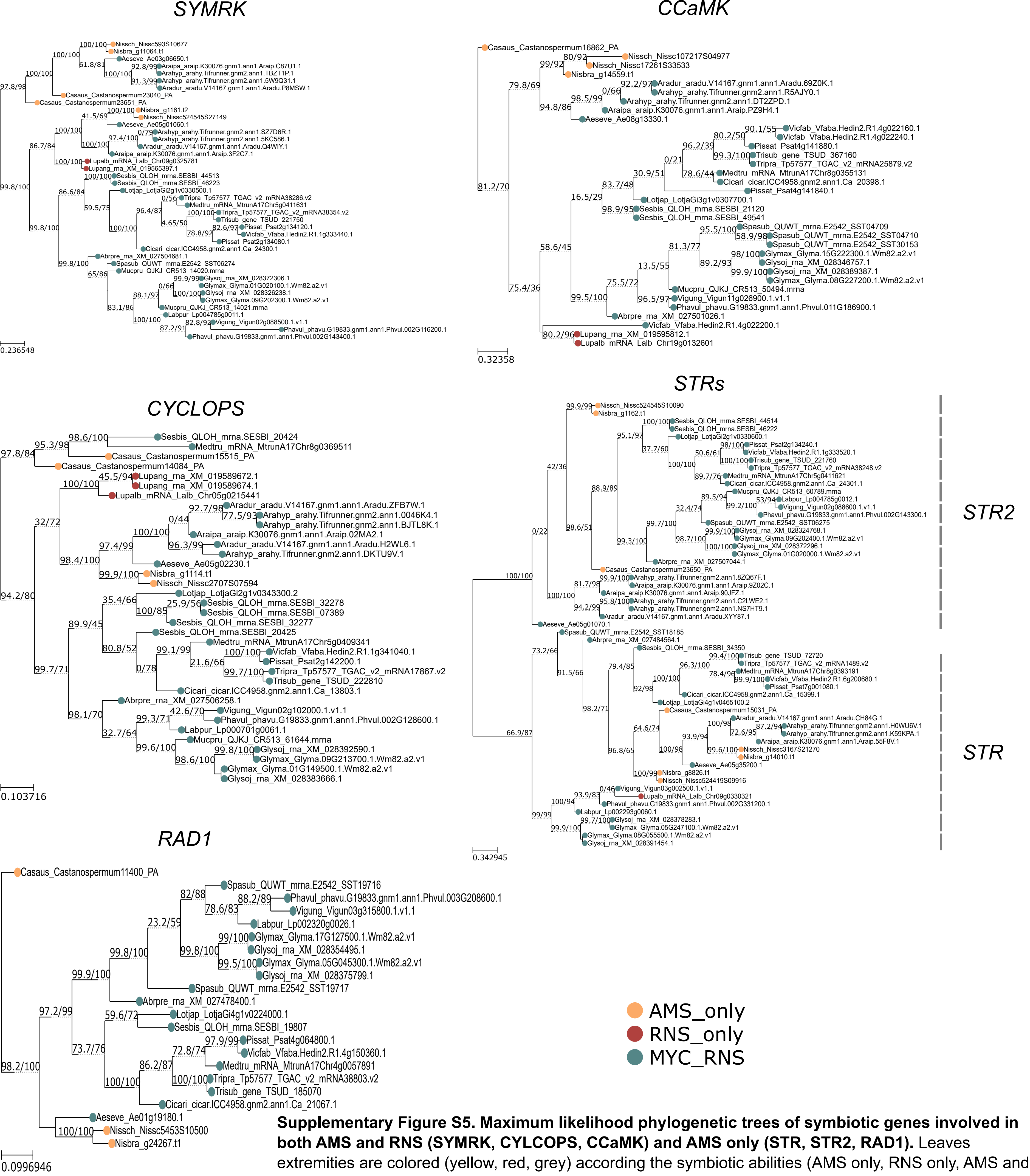


**Supplementary Figure S5. Maximum likelihood phylogenetic trees of symbiotic genes involved in both AMS and RNS (SYMRK, CYLCOPS, CCaMK) and AMS only (STR, STR2, RAD1).** Leaves extremities are colored (yellow, red, grey) according the symbiotic abilities (AMS only, RNS only, AMS and RNS) of each species. Full names of species are available in Supplementary Table S4.
