## Supplementary material for "*Nissolia brasiliensis* as a non-nodulating model legume": Figure S6

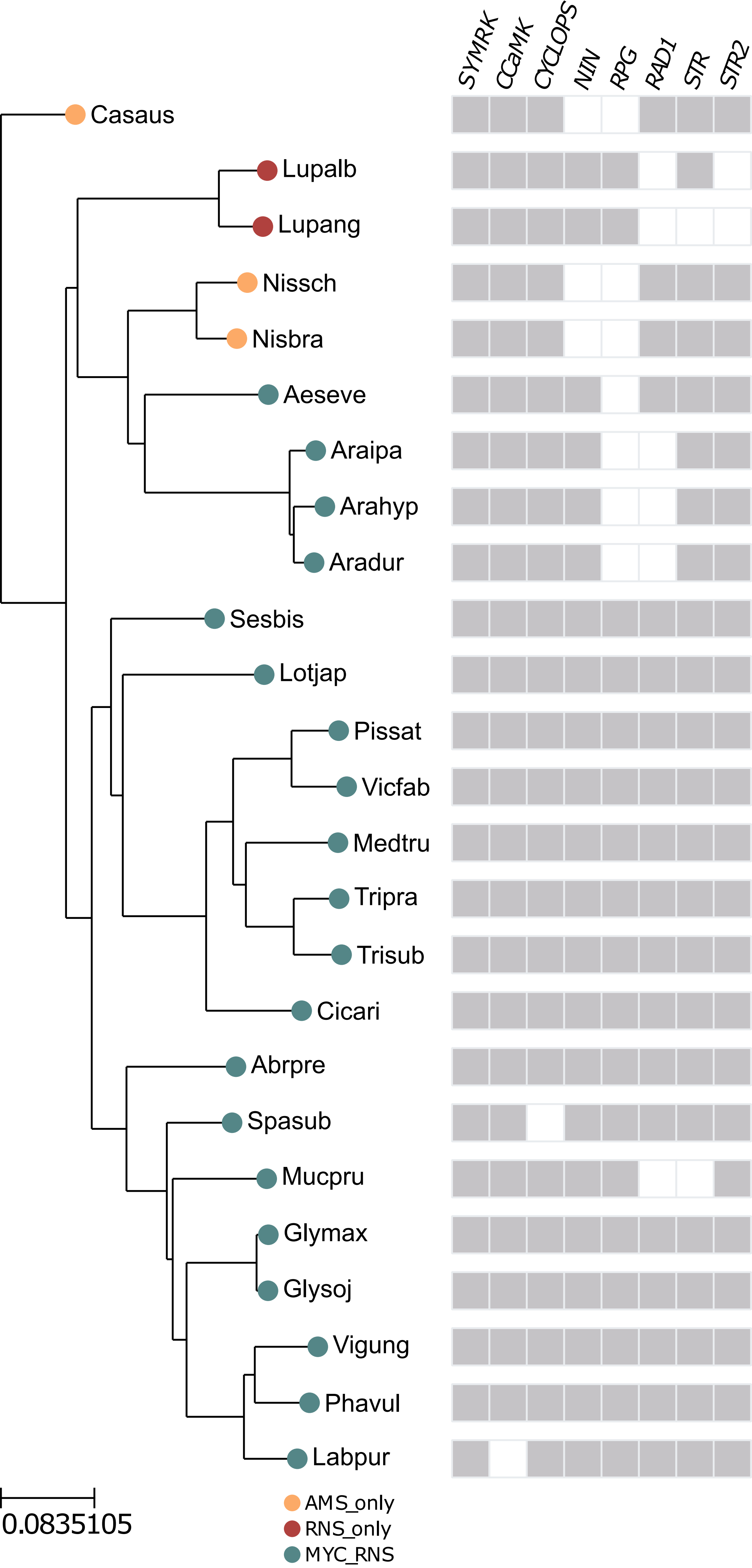


**Supplementary Figure S6. Estimated species tree from OrthoFinder analysis with presence or absence of symbiotic genes** indicated with grey (ortholog present) or empty (ortholog absent) squares. Leaves extremities are colored (yellow, red, grey) according the symbiotic abilities (AMS only, RNS only, AMS and RNS) of each species. Full names of species are available in Supp. Table S4
